# Mitochondrial Membranes: Model Lipid Compositions, Material Properties and the Changing Curvature of Cardiolipin

**DOI:** 10.1101/2023.02.06.527315

**Authors:** Sukanya Konar, Hina Arif, Christoph Allolio

## Abstract

Mammalian and *Drosophila Melanogaster* model mitochondrial membrane compositions are constructed from experimental lipidomics data. Simplified compositions for inner and outer mitochondrial membranes are provided, including an asymmetric inner mitochondrial membrane. We performed atomistic molecular dynamics simulations of these membranes and computed their material properties. When comparing these properties to those obtained by extrapolation from their constituting lipids, we find good overall agreement. Finally, we analyzed the curvature effect of cardiolipin, considering ion concentration effects, oxidation and pH. We draw the conclusion that cardiolipin negative curvature is most likely due to counterion effects, such as cation adsorption, in particular of H_3_O^+^. This oft-neglected effect might account for the puzzling behavior of this lipid.

**SIGNIFICANCE:** Mitochondrial membranes are of fundamental interest to the pathogenesis of neurodegenerative diseases. The biophysics of mitochondrial membranes can be expected to profoundly influence both the electron transport chain and larger-scale mitochondrial morphology. We provide model mitochondrial membrane compositions and examine their mechanic properties. Reconstructing these properties from their constituent lipids, we facilitate the creation of mesoscopic models. Cardiolipin, as the key mitochondrial lipid is given special attention. We find that its mechanical properties, in particular its curvature, are not constant, but highly dependent on specific ion effects, concentration and oxidation state.

## 1. INTRODUCTION

Mitochondrial membranes have a number of unique properties that distinguish them from other cellular and intracellular membranes. At the molecular level, their most prominent features are the almost complete absence of cholesterol and the prevalence of cardiolipin (CL) in the inner mitochondrial membrane (IMM) (1), while the outer mitochondrial membrane (OMM) hosts machinery for lipid and protein transport. Mitochondrial lipids are increasingly studied for their relevance to disease pathogenesis. In particular, neurodegenerative diseases, such as Parkinson’s and Alzheimer’s disease are known to be associated with the oxidation of mitochondrial lipids (2–5) and Barth’s disease is the result of a failure of cardiolipin remodeling (6, 7).

Mitochondria exposed to oxidative stress as well as undergoing other ion and lipid imbalances, exhibit distinct changes in their external and internal morphology (8, 9). Internally, mitochondria contain folded lamellar membrane structures of their inner membranes, the so-called cristae (10). Mitochondrial membrane remodeling is known to be driven by an array of fusion and fission proteins, such as Drp1, the mitofusins, and Opa1 (11), as well as contact sites to the endoplasmic reticulum (12, 13). Yet, these proteins require interaction with mitochondrial lipids to function (14, 15), and the interplay of mitochondrial membrane lipids and the global shape of mitochondria is at present hardly understood. A particular source of confusion continues to be cardiolipin. CL stabilizes the cristae’s structure and functioning and was originally considered a proton conductor (5). This hypothesis is particularly appealing as the lipid also appears in chloroplasts and strongly interacts with the electron transport chain, in particular complex IV. However, it has been shown that at physiological pH, CLs are not protonated (16, 17). At the same time, it is often claimed that CL has a strongly negative curvature and is very fluid. This hypothesis is supported by lipid sorting experiments (18, 19). However, CL is also stable in the lamellar phase (20) – which is seemingly at odds with such behavior and relatively high bending rigidities have been reported for it (21). Perhaps the special features of mitochondrial membranes result from their complex lipid compositions. In experimental setups, typically mixtures of PE, CL and PC lipids, containing a single component for each headgroup are employed as model mitochondrial membranes (22). Therefore, we begin this study with the construction of model mitochondrial bilayers for mammalian mitochondria as well as *Drosophila*. In the next step, we compute the bulk mechanical properties of the resulting systems and compare them to the bulk properties extrapolated from their constituent lipids. For this purpose, we use a recently developed methodology that allows the extraction of bending rigidity as well as tilt moduli and spontaneous curvatures for these lipids (23, 24).

In the next step, we consider the effect of membrane asymmetry, lipid oxidation, curvature and lipid imbalances and the presence of ions. Specific ion-membrane interactions elucidate how the interplay of electrostatics and curvature are the key to understanding mitochondrial membrane dynamics.

## 2. METHODS

### 2.1 Computational Details

Most of the solvated bilayers were constructed using the CHARMM-GUI input generator (25). Typically, the systems contain 200 lipids (i.e. 100 lipids per leaflet) and ca. 10K water molecules. For detailed system compositions, see Table S9. In order to maintain a near physiological salt concentration, 15 mM KCl was added on top of the counterions necessary to neutralize the membranes (where applicable). The lipid types used in this paper are-1-palmitoyl-2-oleoyl-sn-glycero-3-phosphocholine (POPC), 1-palmitoyl-2-oleoyl-sn-glycero-3-phosphoethanolamine (POPE), 1,2-dioleoyl-sn-glycero-3-phos-phocholine (DOPC), 1,2-dioleoyl-sn-glycero-3-phosphoethanolamine (DOPE), 1,2-dipalmitoleoyl-sn-glycero-3-phosphoethanolamine (DYPE), 1,2-dipalmitoleoyl-sn-glycero-3-phosphocholine (DYPC), 1,3-Bis-[1,2-di-(9,12-octadecadienoyl)-sn-glycero-3-phospho]-snglycerol (tetralinoleoyl cardiolipin) (TLCL), 1-stearoyl-2-linoleoyl-sn-glycero-3-phospho-L-serine (SLPS) and L-α-phosphatidylinositol (SLPI). We also refer to the headgroups of these lipids, using the second part of the four-letter abbreviations, e.g. PE refers to phosphoethanolamine. In our simulations, CL is fully deprotonated (−2e charge), as it is known to be at physiological pH values (26). Unless mentioned otherwise, all lipids were modeled using the CHARMM36 force field (27). For one of the CL bilayer system preparations, we used the OPLS-AA force field (28) and in that particular case, we used SPC/E water model (29). Whereas, for all other systems we used the TIP3P model to hydrate the systems (30). Furthermore, when we checked the effect of ionization on the CL bilayer, we replaced half of the K^+^ ions in the system with H_3_0^+^ (31, 32). Simulations were performed at 303.15 K temperature, using the Nosé-Hoover thermostat (33) with a time constant of 1.0 ps. Pressure was maintained at 1 bar using the Parrinello–Rahman barostat (34) in a semi isotropic setup (coupling the membrane plane separately from its normal) with a coupling constant of 5.0 ps. Long-range electrostatic forces were treated using the particle mesh Ewald (PME) method (35) with a real-space cutoff of 1.2 nm. Lennard Jones interactions were cut off at 1.2 nm with a force switching starting at 1.0 nm and a 2 fs timestep was used for integration. Covalent bonds were constrained using the LINCS algorithm (36). All simulations were performed at a constant pressure of 1 bar with a compressibility of 4.5 × 10^−5^ bar^−1^.

After initial relaxation, all the systems were run for 200 ns in an NpT ensemble for lipid bilayer equilibration. For production, the systems were run for 400 ns in an NpT ensemble using GROMACS 2020.3 (37, 38). Bilayer simulations were performed using semi-isotropic pressure coupling. The lateral pressure profile calculation was carried out using the Sega code (39), modified with a Goetz-Lipowsky (40) force decomposition. In order to compute lateral pressure profiles, we ran a separate simulation of 200 ns length with triangular water molecules (as SETTLES are unsupported), which was carried out after pre-equilibration. The neighbor list was updated every 20th step and covalent bond lengths were constrained using the LINCS algorithm. Here, we have increased the LINCS order to 5 in order to deal with the nonlinearity introduced by the constrained H2O ring and turned off force-switching. Pressures and velocities were sampled for 25000 frames to compute the lateral pressure profile.

### 2.2 Extraction of Material Properties

The spontaneous curvature (C_0_), bending modulus (κ_c_) and tilt modulus 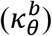, parametrize the Hamm-Kozlov Helfrich (HKH) free energy functional (41–47). These membrane elastic properties are sufficient to model mesoscopic membrane structure, including membrane fusion and fission (48, 49). In addition to the curvature elastic parameters, we also compute the area compressibility modulus *κ_A_*, which is useful e.g. for surface tension computations. Our previously developed computational (ReSIS) approach was used to quantify the κ_c_ and 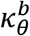 value from local fluctuations of lipid splay and tilt degree of in real space (23, 50–53).

The first moment of the lateral pressure distribution *π*(*z*) is linked to the monolayer spontaneous curvature (*C*_0_) and bending modulus (κ) by (54)

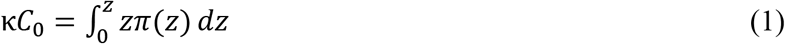

The integration must be carried out along to z axis from the center of the lipid bilayer (z=0) to the bulk, i.e. the integration is carried out over each monolayer separately and then averaged. Using the ReSIS value of κ, we computed the spontaneous curvature (*C*_0_) from Eq. 1. We ensured the membranes to be tension free by requiring the surface tension σto be zero, i.e. subtracting an adequate constant normal pressure from the averaged local, diagonal stress components *σ_aa_*(*z*). We verified the freedom from surface tension by making sure that the required normal pressure, *p_N_*, was close to 1 bar and by checking the z component of the virial in the simulation.

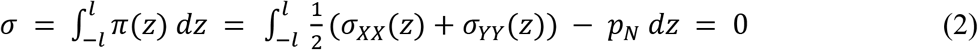

The area compressibility modulus, *κ_A_*, was extracted from the simulations by using

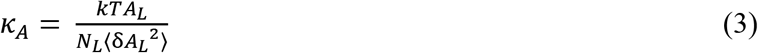

Here, *k* denotes the Boltzmann constant, T is the temperature of the simulation, *A_L_* is the average area per lipid. And δA_L_ is the standard deviation of *A_L_*. This well-known relation (49, 55) is easily derived from the fluctuation-dissipation theorem.

## 3. RESULTS and DISCUSSION

### 3.1. Model Lipid Distribution of Mammalian IMM and OMM

A model lipid distribution for mitochondrial membranes is useful for experiments and MD simulations, e.g. of mitochondrial membrane proteins. In order to build a realistic mitochondrial membrane model, we considered both the acyl chain composition of mitochondrial membranes and their head group composition. We follow Daum’s example by using rat liver mitochondrial headgroup composition to represent mammalian cells (56). Details and literature are given in Table S1 and S4. The composition of acyl chains for lipids in living organisms is very diverse; nevertheless, there are usually only a few species that dominate the fatty acid composition of each lipid. Thus, the number of species was limited in a pragmatic manner to few, commercially available lipids. The data by Daum does not contain any assignment of the acyl chains to lipid headgroups. Assigning these to the tails is nontrivial. We departed from the membrane specific acyl chain determination by Ardail et al. (57). We consider the length of acyl chains to be less important to the lipid physical properties than lipid (un)saturation. Accordingly, we simplified the total acyl chain composition based on the saturation type, interpolating between data by Daum (58) and lipidomics data by Oemer et al. (59). Arachidonic acid (C20:4) and linoleic acid (C18:2) are grouped as unsaturated lipid type. It is known that 18:2 CL is dominant in rat liver (60). Hence, we chose this as the exclusive CL tail. Due to our simplification of 20:4 to 18:2 tails, we chose 16:0/18:2 for PI and 18:0/18:2 for PS lipids, with good experimental support from rodent mitochondria, in the sense that there is a dominant species with a saturated and a polyunsaturated tail for both PS and PI (61, 62). We note, that arachidonic acid appears as the dominant unsaturated species, especially for PI, but expect the effect on bulk properties to be minor.

Further details on our simplification are given in Table S2 and S3. The distribution for PC seems broad; we choose two variants of PC, 1-Palmitoyl-2-oleoyl (C16:0/18:1) and di-palmitoyl (C16:1/16:1) (62). As a result, our final composition has a lower 18:0, and higher 18:1 content than the experimental data. The resulting, slightly lower fraction of fully saturated chains is necessary to keep all components of the membrane fluid at the simulation temperature, for separate mechanical evaluation. For PE, we chose POPE to fulfill the total (simplified) composition with a typical acyl configuration. The simplified acyl chain for the OMM was constructed in an analogous manner. Here, we considered two acyl variants (16:0/18:1 and 16:1/16:1) for PC and PE, a single variant for PI (16:0/18:2), CL (18:2/18:2), and PS (18:0/18:2). We then estimate each acyl chain variant which is distributed among each phospholipid, as done before. The calculated percentages of acyl chain variants are shown in Table S3. The final model lipid compositions in mammals based on acyl chain and head groups for IMM and OMM are shown in Fig. 1.

**Figure 1:**
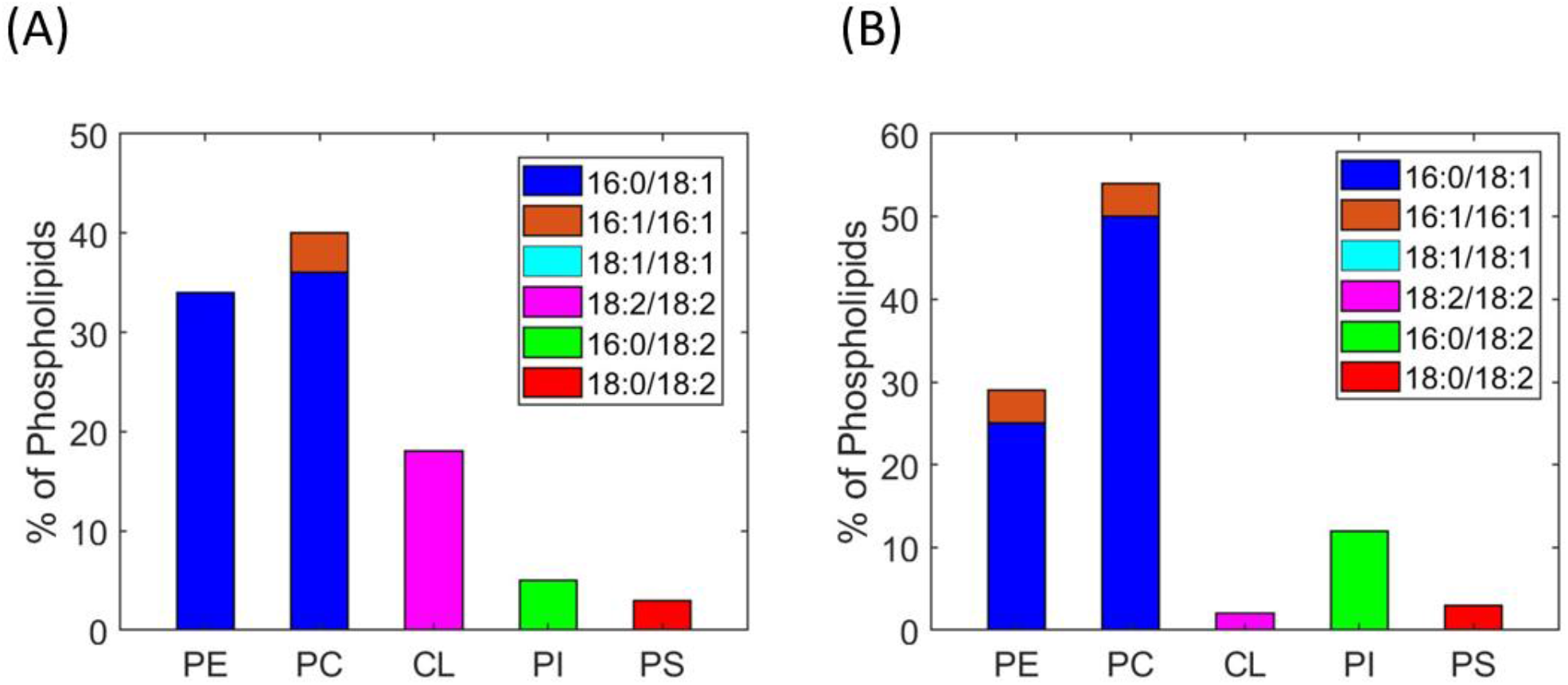
Model lipid compositions of mammalian mitochondrial membranes. Lipid distribution in mammalian (A) IMM and (B) OMM showing head group as well as fatty acyl chain of the lipid.

Mammalian mitochondrial membranes have more PC content in both IMM and OMM than PE. However, as is well-established, CL content is far higher in the IMM than in the OMM.

### 3.2. Construction of IMM and OMM Model Lipid Compositions for *Drosophila*

Acehan *et al*. (63) elucidated the total headgroup composition of *Drosophila* mitochondrial membranes. However, the compositions of IMM and OMM were not measured separately. Hence, we started by creating a model of *Drosophila* mitochondrial membranes in combination with other data. The first step was to estimate the distribution of these headgroups across the mitochondrial membranes.

The percentage of phospholipid headgroups in IMM and OMM of mammals is known from the literature and given in Table S1. In order to build a membrane model for *Drosophila*, we need to estimate the ratio of the number of lipid headgroups in IMM and OMM (N_IMM_ / N_OMM_). Furthermore, we determine the lipid fraction in IMM (χ_IMM_)and OMM (χ_OMM_). Then, we estimate the average total percentage, or, in our case of 100 lipids per leaflet, the number of phospholipids 〈N_*Lipid*_〉 for each leaflet as follows:

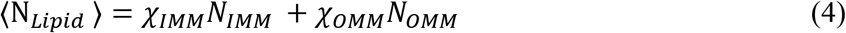

Where, χ_IMM_ and χ_OMM_ are the fraction of lipid in the inner and outer membrane, respectively. We take these fractions to be 0.8 and 0.2, meaning that 80 % of total lipids are in the inner mitochondrial membrane. Knowing the total lipid composition from literature and taking the headgroup distribution ratio to be equal to that found in mammals (i.e. estimating the ratio of, e.g. PE in inner vs outer membrane to be conserved), it becomes possible to estimate the headgroup composition for OMM and IMM respectively.

We controlled our estimate of χ_IMM_ and χ_OMM_, by reproducing the average headgroup composition of phospholipids from the data of Horvath and Daum (56). Furthermore, we find that our estimate also reproduces the total lipid compositions of plants well (for which total and relative data are given by Daum). The resulting headgroup composition is shown in Table S5.

### 3.3. Full lipid distribution of IMM and OMM for *Drosophila* by combining acyl chain and head group compositions

We obtained the acyl chain composition from Dubessay *et al*. (64) and used it to estimate the lipid distribution of the IMM for Drosophila. Following the simplification and head assignment procedure we described for mammals in section 3.2, we considered three acyl variants for PC, PE and a single variant for the charged PI, CL, and PS lipids. These were based on the lipid simplification (grouped according to lipid saturation/unsaturation) given in Table S6 and S7. After assigning PI, CL and PS as before, we distributed the rest of the acyl chains into the neutral lipids equally into the IMM and OMM to fulfill the total acyl chain composition requirement given in Table S5. Results are shown in Fig. 2.

**Figure 2:**
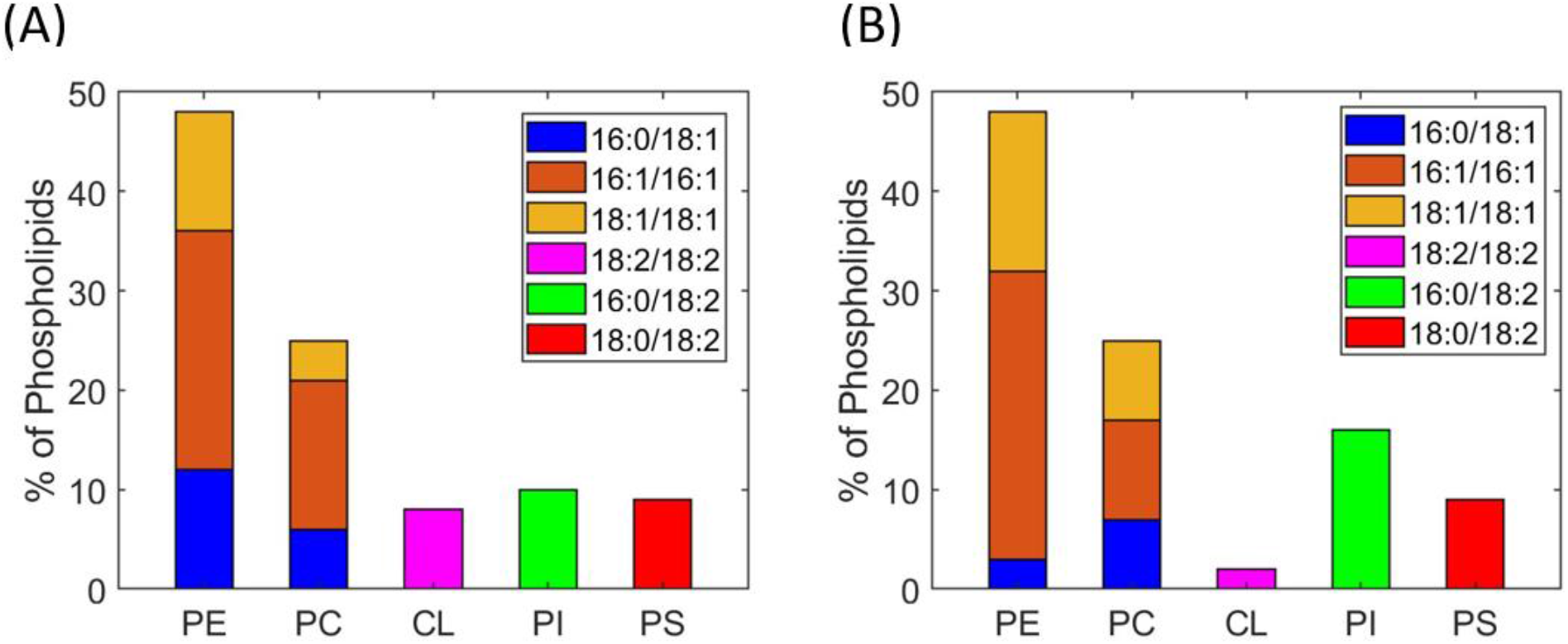
Model lipid composition of *Drosophila* IMM and OMM. Lipid distribution in (A) IMM and (B) OMM showing head group as well acyl chains of the lipids.

It follows from our approach that due to the higher PE content found in the mitochondria, both in IMM and OMM, PE is the dominant head group. Accordingly, the main differences between IMM and OMM arise from their CL and PI lipid content. CL is only present in IMM, whereas PI is typical for the OMM.

### 3.4. Model of the asymmetric lipid distribution of the mammalian IMM

Lipids are asymmetrically distributed in the inner and outer leaflets of biological membranes. In particular, CL is localized mostly in the mitochondrial matrix side of the IMM. An asymmetric membrane lipid distribution between outer and inner leaflets might show some significant changes in their biophysical properties like bending, rigidity and spontaneous curvature for each monolayer. We selected the mammalian system as there is a known lipid headgroup distribution for the inner and outer leaflets of IMM and OMM (56). Area mismatch between two asymmetrical leaflets can lead to one of the leaflets being under tension, whereas another might be compressed. In order to avoid this, we simulated each membrane leaflet in a separate, symmetrical configuration. We verified that the total area difference was small (see Fig. S2) and then we assembled the asymmetrical mitochondrial membrane by combining the two leaflets. In our asymmetric membrane model, CL and PI are dominant in leaflet facing the matrix and other lipids are distributed in both leaflets. The lipid distribution in the asymmetrical membrane model was obtained by sorting the mammalian IMM model composition according to (56). However, the data in that reference does not really admit to matching the areas, due to the area per lipid (APL) of CL being ca. twice that of other lipids. Accordingly, we distributed some PC and PS into the CL depleted intermembrane space (IMS) side, see Table S8 for full details. Our proposed asymmetrical IMM model composition for mammals is shown in Fig. 3.

**Figure 3:**
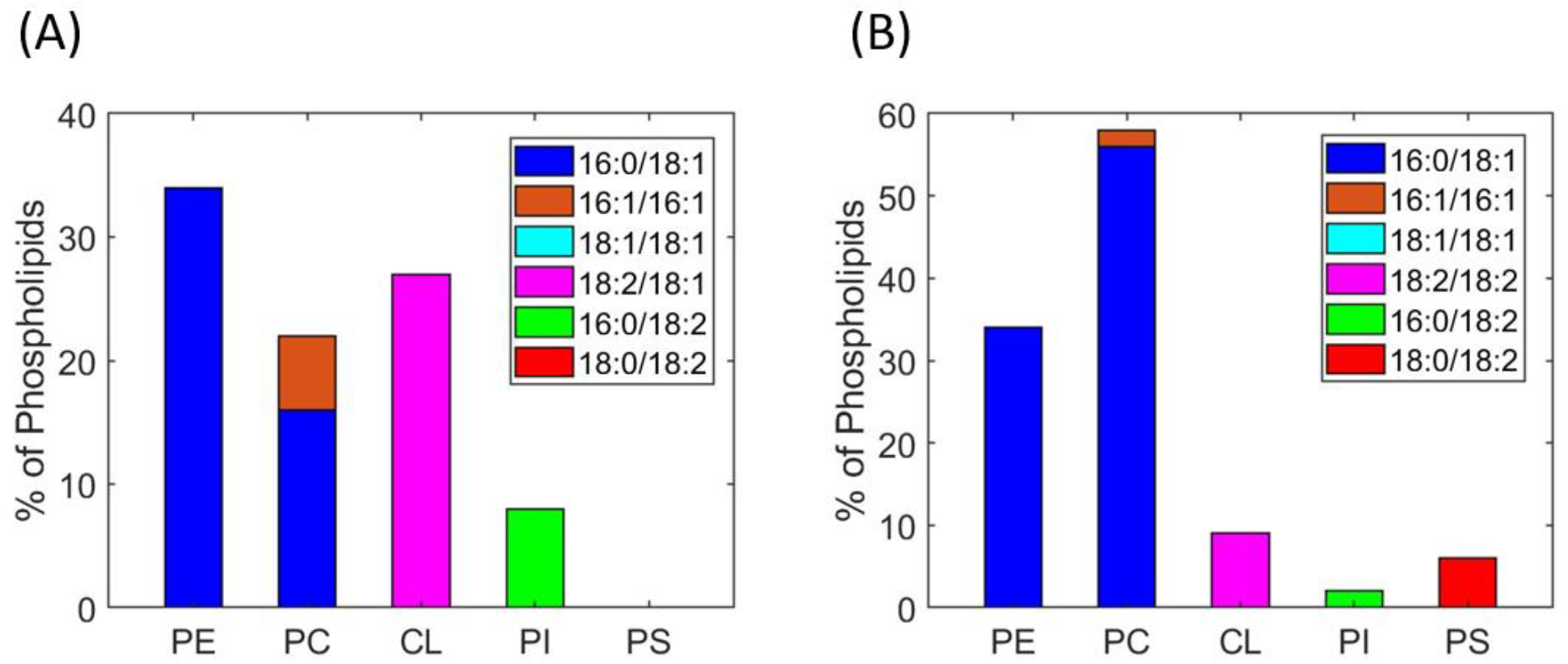
Asymmetric lipid distribution of inner (matrix) and outer (IMS) leaflets of mammalian IMM. Lipid distribution in inner leaflet (A), outer leaflet (B).

### Material Properties of the Membranes

### 3.5. Area per lipid and thickness of the lipid bilayers

To understand the structural properties of the bilayers, we first computed the APL and lipid bilayer thickness of the model mitochondrial membranes (IMM and OMM) for both symmetric and asymmetric membranes as listed in Table S11. In addition to the model mitochondrial membranes, we also computed the properties for single component lipid membranes for all their components, with the ultimate goal of checking the validity of commonly used mixing rules for membranes. APL and bilayer thickness of these lipid variants are also listed in Table S10. The APLs and bilayer thicknesses of single component DOPC, POPC, DOPE and POPE bilayer systems are in agreement with previously reported data (49, 65, 66).

We will now focus on cardiolipin (CL). Because of the relevance of this lipid, we computed its structural and elastic properties using different forcefields and system geometries, namely CHARMM36 and OPLS-AA. The APL for CL [(18:2)_4_] system using the CHARMM36 force field is 134.29 Å^2^, which is slightly higher than using the OPLS-AA force field (132.33 Å^2^). The CL-OPLS-AA force field shows better agreement with the reported experimental data (129.8 Å^2^) by Pan *et al*. for TOCL [(18:1)_2_:(18:1)_2_] (67). Cardiolipin oxidation is considered an important potential factor for mitochondrial diseases. Therefore, to examine the potential changes due to CL oxidation, we have also considered an oxidized form of cardiolipin (CLox) by substituting one of the acyl chains with 13-hydroperoxy-trans-11, cis-9-octadecadienoic acid (13-tc) in OPLS-AA (68). CL_ox_ has higher APL (~140.209 Å^2^) than CL (~132.334 Å^2^) using OPLS-AA. Note that this is far from the only possible oxidation product, but simulating single-component membranes composed of more oxidized forms, such as lysolipids will destabilize the bilayer too much.

As a first elastic property of membrane bilayers, we have calculated the area compressibility (bulk) modulus (κ_A_). Results are given in Figure 4 (A, B). Values for DOPC and POPE match quite well with those reported by Lee *et al*. (25). For the POPC bilayer, the κ_A_ value is 182.08 pN/nm. This value is significantly different from the value reported by Lee *et al*. (25). The difference is ~ 100 pN/nm. However, Piggot *et al*. reported that the κ_A_ value varies from 180-330 mN/m for POPC (69). Thus, we can conclude that this difference is within the margin of error.

**Figure 4.**
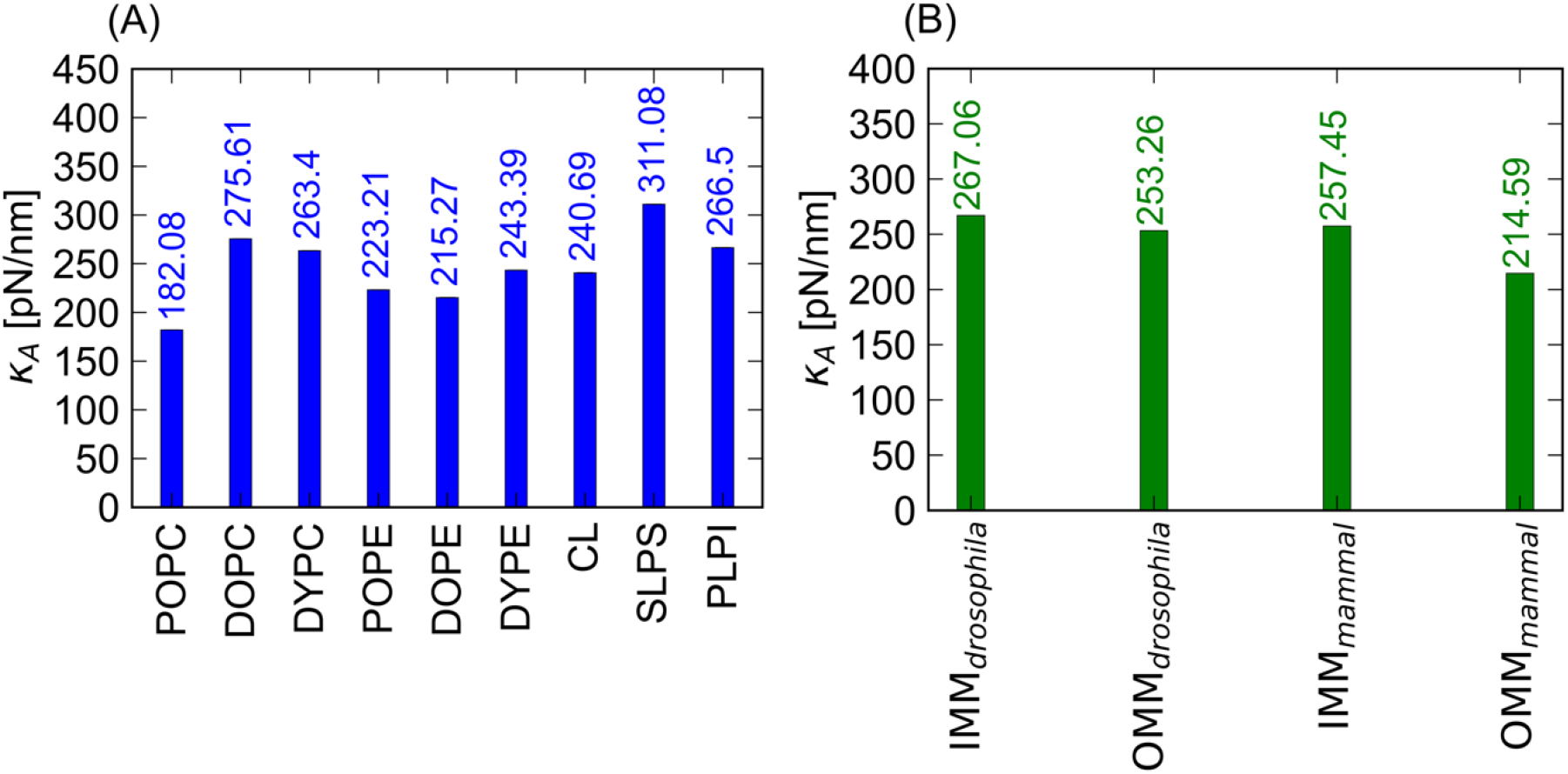
Area compressibility moduli of the different lipid bilayers. Compressibility moduli (Kappa) (in pN/nm) of (A) the bulk systems and (B) the mesoscopic model of mitochondrial membranes for drosophila and mammals.

Pan *et al*. reported that charged lipid bilayers (such as POPS) possess larger κ_A_ values than neutral lipid bilayers (such as POPC) (70). Thus, we would expect that charged lipid bilayer systems have larger κ_A_ values as compared to neutral lipid bilayer systems. The κ_A_ values (in Figure 4 A) clearly suggest that SLPS has a larger area compressibility modulus than other single lipid systems.

Figure 4 B shows the κ_A_ value for the mitochondrial membrane for mammals and *Drosophila*. The Kappa value generally varies from 214-300 pN/nm. We find that the κ_A_ values of single component membranes are similar to these of the mitochondrial model membranes. It appears that there is no general trend for either the IMM or the OMM to have larger or smaller values, and there seems to be no definitive statement possible on the differences between species, with the potential exception of the low area modulus for the mammalian OMM.

### 3.6 Monolayer Bending Rigidities, Spontaneous Curvatures and Tilt Moduli

The bending modulus κ_c_ and the spontaneous curvature C_0_ are some of the most important parameters that determine the mesoscopic behavior of membranes, as they describe the resistance against bending deformations and their preferred geometry. If we want our mitochondrial membrane models to be available for mesoscopic modeling studies, we need to determine the membrane elastic properties under different conditions, including mixtures. We want to address the question – ‘will the different mitochondrial membrane lipid composition across species affect the bulk properties of the membrane or not?’ While different types of ions (divalent and monovalent) modulate the membrane elasticity (71–75), we probe the case of the noncoordinating salt KCl. The lipid composition of IMM/OMM for *Drosophila* significantly differs from the lipid composition for mammals. In order to find out whether this difference leads to important changes in the elasticity of the membrane, we calculated κ_c_, C_0_ and the tilt modulus, 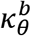 for the model membrane sytems (shown in Fig. 5 C and D, Fig. S1 B). Simultaneously, we calculated the κ_c_ and 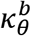 for the bulk lipid systems (Fig. 5 A and B, Fig. S1 A).

**Figure 5.**
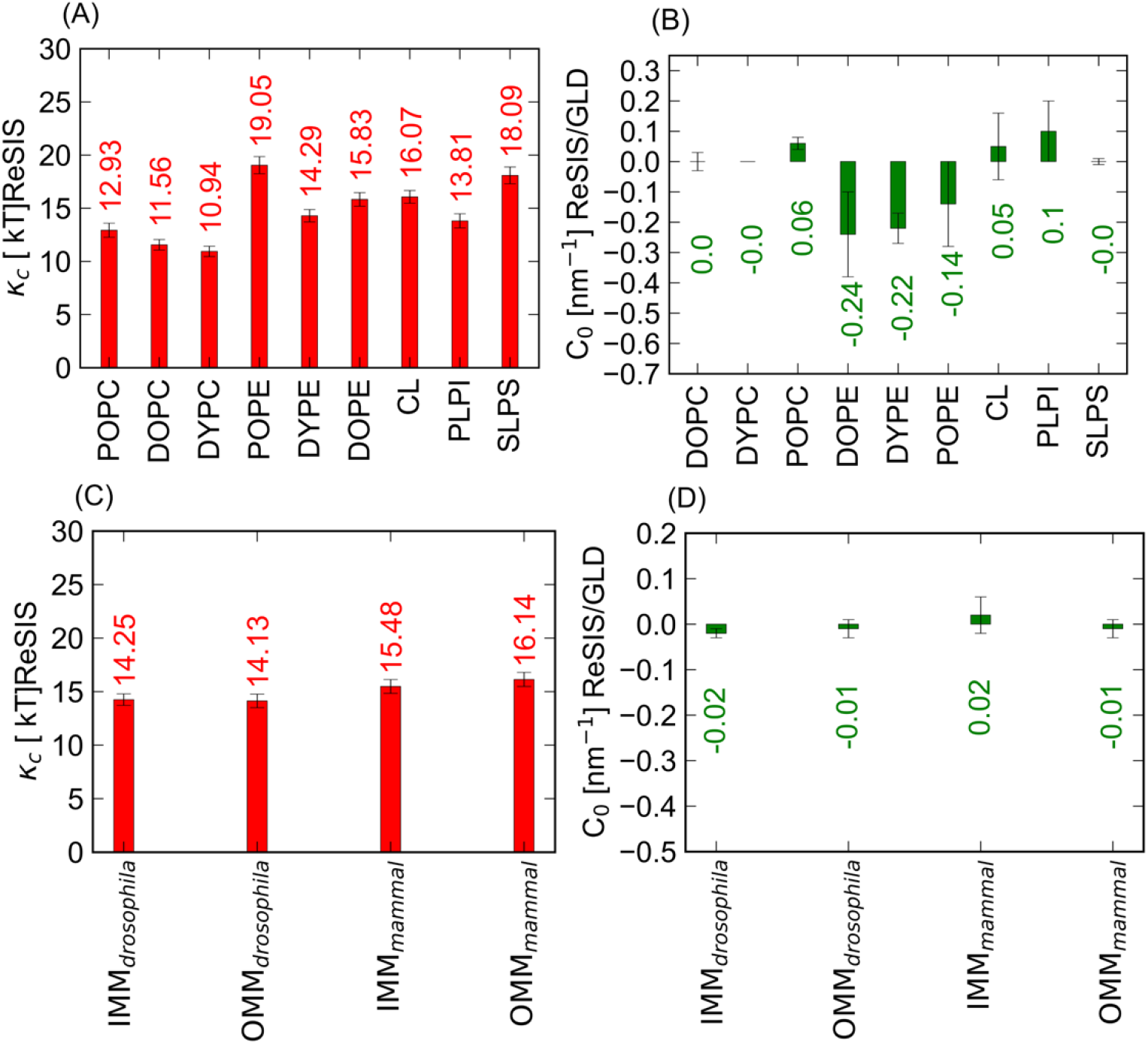
Elastic properties and spontaneous curvature of the bulk system and model mitochondrial membranes (IMM and OMM) for both species. (A) Monolayer bending rigidities (κ_c_) (in kT) and (B) Spontaneous curvature (C_0_ value in nm^−1^) of bulk systems (C) Monolayer bending rigidities (κ_c_) (in kT) and (D) Spontaneous curvature (C_0_ value in nm^−1^) of IMM and OMM of *Drosophila* and mammal. Error bars show the ΔC_0_ value (it is calculated based on the difference of the C_0_ values of each monolayer)

Although the IMM and OMM compositions of the model membranes (*Drosophila* and mammal) are different from each other, their κ_c_ and 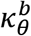 values do not differ significantly. In contrast to this, large differences in all of these parameters occur for the single-component lipids. We do not know whether the homogeneity of bending elastic parameters is a coincidence or it reflects the diversity of evolutional outcomes, which lead to functional membranes.

C_0_ is an important parameter as well as lipid-specific property, hence we computed the C_0_ values for our model mitochondrial membrane. Figure 6 shows the C_0_ values for IMM and OMM for two types of species and the asymmetric membrane. Although, the IMM compositions of *Drosophila* and mammals differ significantly, their C_0_ values do not. Both are ca. 0 nm^−1^. In conclusion, we observed no significant difference in the bending rigidity and spontaneous curvature between these two models. This stands in marked contrast to the behavior of the single component lipids, which show marked differences in spontaneous curvature, as is expected (24, 49). The material properties suggest that the unique biological properties of mitochondrial membranes are not due to *global* curvature elastic terms but might significantly change via *local* demixing induced, e.g. by proteins. Another potential issue is whether the curvature elastic properties of mitochondrial membranes are significantly affected by their asymmetric lipid composition. Several examples of the importance of membrane asymmetry suggest that important differences are possible, especially for domain forming lipids (76–78). Figure 6 (A, B) shows the bending rigidities (κ_c_) and spontaneous curvatures (C_0_) for the asymmetric membranes. Initially, we found that κ_c_ of IMM (mammals) and asymmetric type IMM (IMM_asymm_) is almost the same, i.e. 15.4 and 15.7 kT. We calculated κ_c_ for IMM and separately for both leaflets (inner and outer leaflets) of the IMM. Here, IMS_a_ is the outer leaflet and Matrix_a_ is the inner leaflet of asymmetric IMM, where inner refers to the mitochondrial matrix side, and outer leaflet represents part of the membrane facing the intramembrane space side. Surprisingly, we did not observe any significant differences between the κ_c_ for each leaflets of the IMM_asymm_. We further examined asymmetry and possible interdigitation effects by comparing the monolayer κ_c_ values Matrix_a_ and IMS_a_ to properties computed from a symmetric bilayer of the inner and outer IMM composition. These are marked with IMM_b_ and do not differ significantly from the monolayer values. Similarly, the C_0_ for Matrix_b_ and IMS_b_ are within the margin of error for the C_0_ for IMM (mammals) and asymmetric IMM. This result clearly suggests that for the purpose of global material properties of mitochondrial membranes, asymmetry is not a strong factor. Interestingly, the C_0_ for the leaflets Matrix_a_ and IMS_a_ are not those of their respective symmetric analogues, but follow the slight membrane compression and stretching from due to the very small area mismatch. Following Deserno, we suggest that the experimentally observed differences in for asymmetric membranes might occur due to packing differences related to differential stress (77).

**Figure 6.**
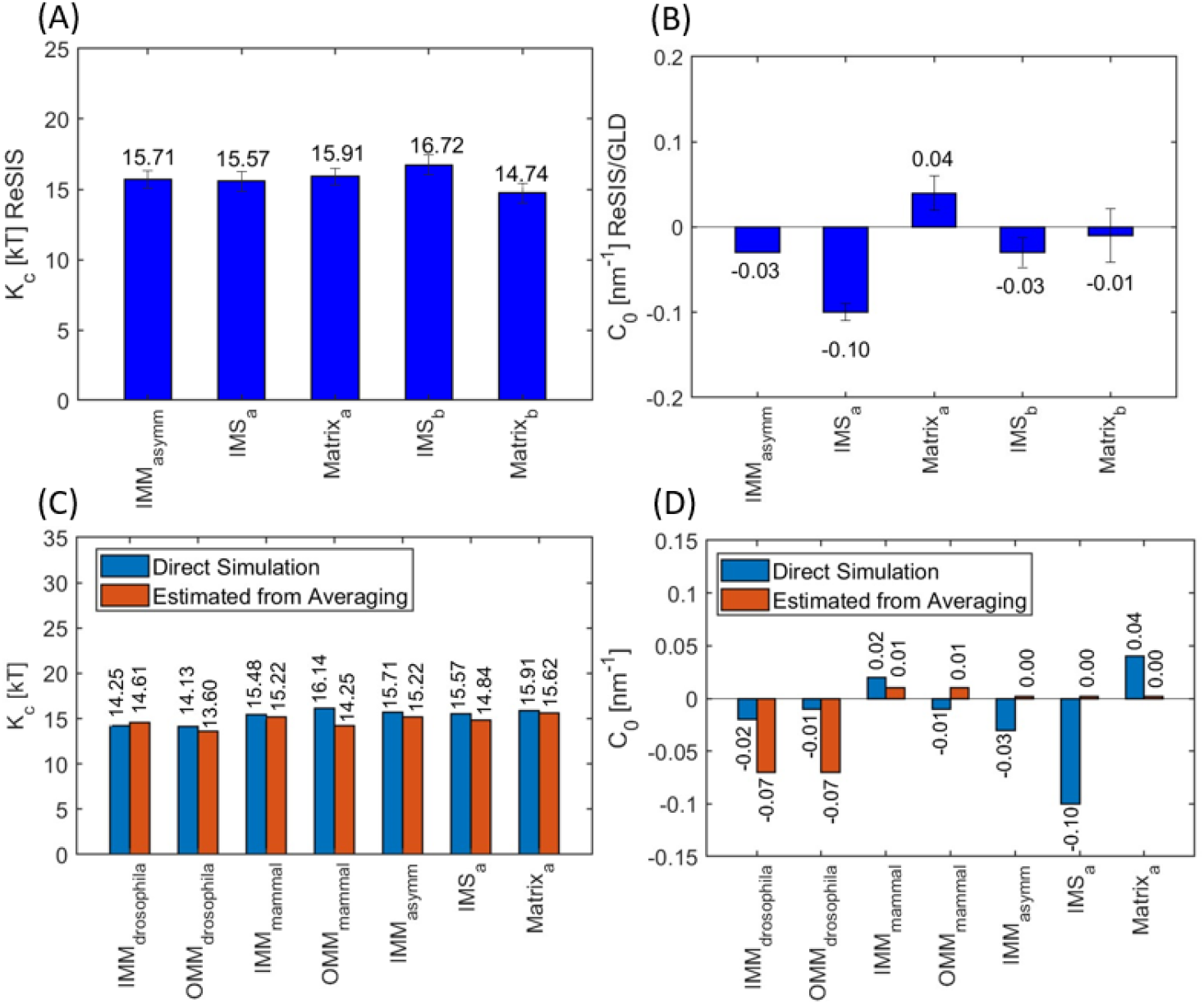
Comparison of the elastic properties of the asymmetrical model IMM. (A) bending rigidity (κ_c_) (in kT) (B) Spontaneous curvature (C_0_ in nm^−1^) of IMM for mammals, asymmetric membrane, inner and outer leaflet of IMM, respectively. (C) Monolayer bending rigidity (κ_c_) (in kT) of mitochondrial membranes of *Drosophila* and mammals using ReSIS methodology and the direct simulation method. (B) Spontaneous curvature (C_0_ in nm^−1^) of the corresponding membranes.

### 3.7 Extrapolation of Elastic Properties of IMM and OMM by Simple Mixing Rules

Just like any other biological membrane, mitochondrial membranes are locally inhomogeneous. CL is known to be strongly sorted by negative curvature in experiments (18, 19). This stands in direct, apparent contradiction to our results which show negligible curvature. To make a useful mesoscopic model without re-extracting properties for any composition, we need a way to generate the elastic properties for different compositions. Our simulation of all the lipid components in one component system and the consistent extraction of their simulated properties enables us to test well established empirical equations for the mixing of membrane properties. As proposed (53), the bending rigidity of a lipid mixture is estimated via a harmonic mean of the corresponding values of its constituents. Accordingly, the bending rigidity (*κ_c_*) of the mitochondrial membrane can be calculated using,

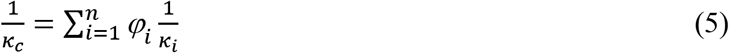

where, *φ_i_* represents the area fraction of the individual lipid component (PC, PE, CL) present in the mitochondrial membrane. The index *i* runs over the individual lipid types present in the IMM or OMM. The *κ_c_* values for each bulk lipid system are shown in Fig. 5A. Likewise, the spontaneous curvatures (*C*_0_) of different IMM and OMM compositions are typically estimated in the literature (79, 80) by direct averaging over the spontaneous curvatures of the constituent lipids (here the areas per lipid are neglected, see Fig. 5B for the values used):

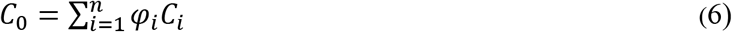

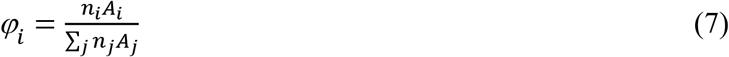

Here, *φ_i_* represents the area fraction, *A_i_* refers to an area per lipid (APL) of the lipid species *i* and *n_i_* refers to the number of lipids of type present in the bilayer. Additionally, it is considered that the APL does not change during lipid mixing.

Figure 6 C shows the estimated values for *κ_c_* obtained by using these averaging procedures. The predicted average value of *κ_c_* matches well with the ReSIS method for both species of mitochondrial membrane (IMM and OMM) (Fig. 6A). This is a very encouraging result, as it allows us to simultaneously confirm the consistency of the ReSIS results and explain the very similar properties found for the different types of mitochondrial membranes.

Figure 6 B shows the *C*_0_ values for IMM and OMM of *Drosophila* and mammals as estimated by Eq. 6. Here, the predicted average value of spontaneous curvature shows some differences with the values obtained by the direct simulation method (Figure 6B). However, this difference is not very large. Thus, we conclude that these two equations are helpful in predicting the elastic nature of the membrane when its composition is known. The estimation from the components allows us to construct curvature elastic properties for any realistic mitochondrial membrane headgroup composition. That might be of value in future, more detailed mesoscopic models.

We have noted that the predicted curvature from the components somewhat deviates from the curvature found in direct simulation. The deviation is definitely not large (with respect to the method errors), but it may point towards some potential sources of disagreement. For example, we have not considered the effect of counterion distribution and the reduced charge density when going, e.g. from single-component CL or PS to a membrane mixture. Overall, it is surprising that the combined deviation seems this small, considering the rough approximations made.

### 3.9 The Puzzle of Cardiolipin

Our extraction of mitochondrial membrane material properties has not resulted in any special properties for CL; we have not obtained a low spontaneous curvature for CL. Boyd *et al*. already predicted a very small absolute C_0_ value of bulk CL system (6). Conventional wisdom suggests that due to its small head group and relatively large tail (four acyl chains), CL must have a negative intrinsic curvature. The main empirical support for an extremely low spontaneous curvature of CL stems from lipid sorting experiments. It was found that CL sorts towards the inner leaflet of LUVs and membrane tubes with a high degree of selectivity (18, 19). Assuming ideal mixing entropy for CL, this fact can only be explained by extremely high negative values for C_0_. However, there are good reasons to assume that such strong negative curvature is unrealistic. For example, it is possible to create lamellar phases made of pure CL. It is of particular interest to mitochondrial pathologies to also check the effect of lipid oxidation for CL.

The mono-peroxide form of CL has a lower κ_c_ (12.04 kT) than the native form (16.98 kT), which can be explained by the less favorable lipid tail packing and the concomitant increase in APL (~140 Å^2^). Overall, the tilt modulus of all CL variants is low in comparison with its bending rigidity (Fig. 7C). This may have profound structural implications, e.g., by privileging tilt deformations over bending deformations. It might also explain the widely held view that CL is a “soft” lipid. The apparent softness of CL would then be due to its chain tilt and not due to its susceptibility to bending deformations.

**Figure 7.**
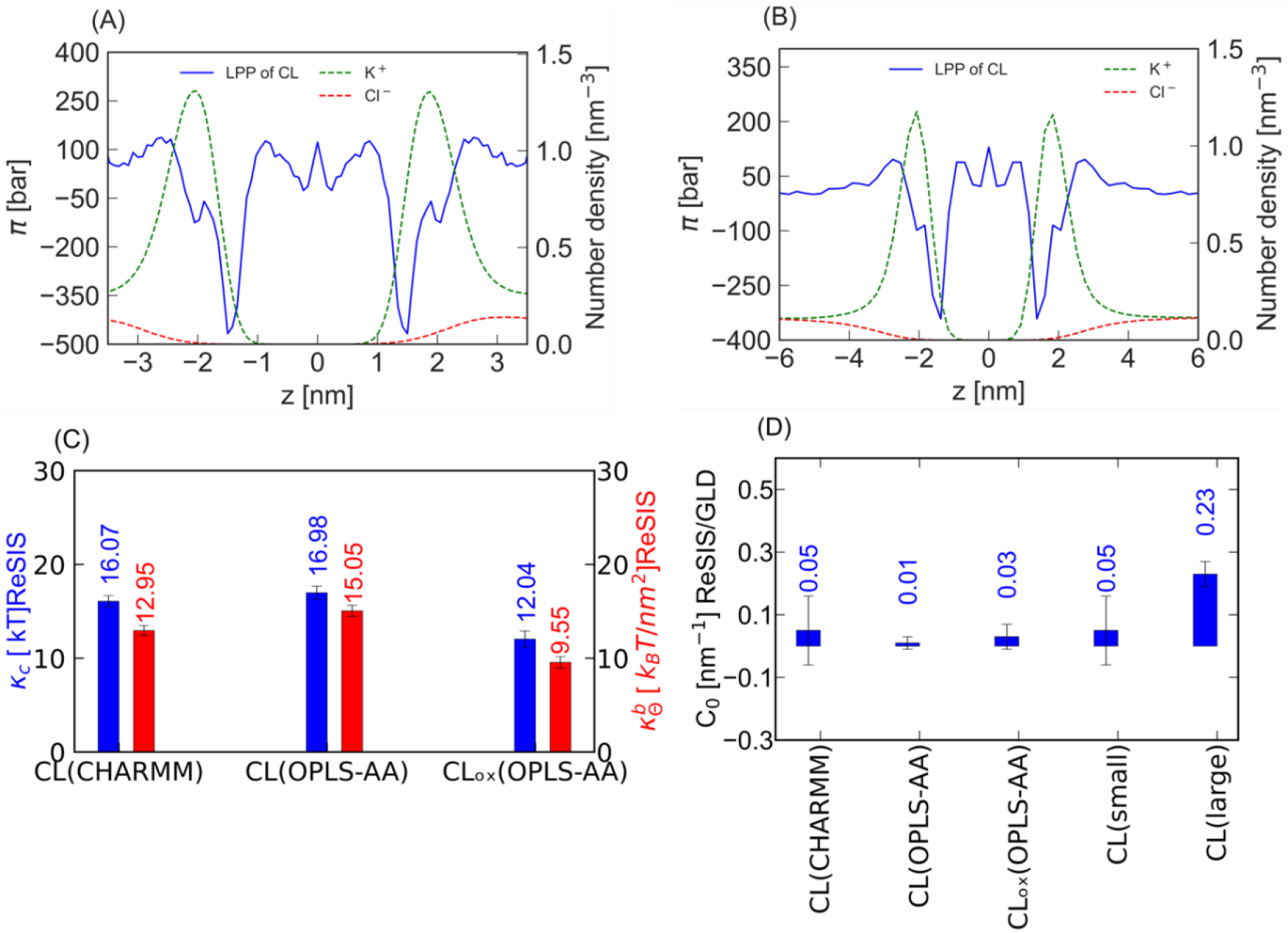
Lateral pressure profile of CL for different simulation box sizes. Lateral pressure profile of CL in the presence of (A) small and (B) large number of H_2_O, with K^+^ and Cl^−^ ion distribution along the bilayer normal. (C) **Elastic properties of CL using different force fields.** Bending rigidity (κ_c_) (in kT) and tilt modulus 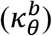 (in kT/nm^2^ unit) of Cardiolipin (CL) for different force fields as well as normal and oxidized forms. In this figure, CL(OPLS-AA) represents the normal form of CL, whereas CL_ox_(OPLS-AA) represents the oxidized form of CL using the OPLS-AA force field. The error bar shows standard errors. (D) **Spontaneous curvature of CL for different force fields.** The spontaneous curvature (C_0_ in nm^−1^) of CL is given at different hydration levels, force fields (CHARMM and OPLS-AA) and in its oxidized form. In this figure, CLox(OPLS-AA) represents the oxidized form of CL using the OPLS-AA force field.

The ability of CL to form lamellar phases is highly dependent on the salt concentration (20, 81). Recently, Allolio and Harries proposed that a long-range lipid clustering by divalent Ca^2+^ ion induces a negative curvature formation. In this study, we discovered that the distribution of the counterions will also affect the spontaneous curvature of lipids, independently of them being bound to headgroups or not. The stress profile of a lipid affects the curvature according to the first moment of the pressure tensor (see Eq. 1). Accordingly, the same contribution of an ion to the stress will lead to a more positive contribution to C_0_ if it occurs further away from the bilayer center. Due to screening, an increased salt concentration leads to a decreased ion-ion repulsion, as expressed by the Debye length. Thus, salt concentration might play a very important role in regulating the intrinsic curvature of CL. This is well established experimentally, as CL can be transferred from a lamellar phase into an inverted hexagonal phase by increasing the salt concentration (82).

Thus, both salt concentration and the distance of the counterion density influence the curvature of CL. In Figures 7A and 7B, we show the lateral pressure profiles of a CL bilayer at different repeat spacings of the simulation. We immediately notice the very strong negative lateral stress contribution of the CL headgroup region. This finding confirms the geometric reasoning based on the small headgroup size. However, the contribution of the counterion density offsets the negative pressure of the headgroup, by its levered positive contributions via the Maxwell (and osmotic) stress contributions. The larger the d-spacing, the more this effect would be expected to contribute to the curvature.

Integrating the pressure profiles confirms the intuition obtained from inspecting the lateral stress profiles. As shown in Fig. 7D, the more dispersed counterion density leads to a larger value for C_0_. The balance between ion dissociation, CL electrostatics and curvature might be highly forcefield dependent and, therefore, in Fig. 7C, the κ_c_ value of CL in both CHARMM and OPLS-AA force fields are reported. They are found to be similar. For identical box sizes, the values of C_0_ are also comparable. This allows us to rationalize the stability of the lamellar phase of CL at low ion concentrations.

It thus becomes evident that the integration of CL into any sort of mesoscopic model necessitates the incorporation of electrostatic terms generated by the counterion profile. It also raises obvious questions with respect to the validity of coarse-grained models of CL without correct long-range electrostatics and water polarization. In order to check for ion-specific effects, we replaced half of the K^+^ ions with the H_3_O^+^ ions and repeated the MD simulation of the single component CL system. In Figure 8 A, the C_*0*_ values of CL in the presence of K^+^ ions and both K^+^ and H_3_O^+^ ions are shown. The data shows that the addition of H_3_O^+^ ions in the bulk CL system induces negative spontaneous curvature in the bilayer. Figure 8 B shows the lateral pressure profile of CL and the number density of K^+^ and H_3_O^+^ions. Our data shows that the head group region of CL is strongly enriched in the H_3_O^+^ ions over the K^+^. Thus, we propose that the H_3_O^+^ ion shows an ion specific interaction with the head group (phosphate group) of CL. And we believe this kind of interaction leads to negative spontaneous curvature in the bilayer. The reason for this induction of negative curvature is to be found both in the depletion of ions at a large distance to the bilayer center and in the screening and binding effects of the H_3_O^+^ ions at the membrane interface. At this point, it should be noted that we have not protonated the CL lipids but merely have put them in contact with water self-ions. Therefore, it is expected that no sharp transition should be visible in a titration curve, i.e. that lipid properties gradually change in a qualitative manner before the protonation of CL is achieved. An enhanced presence of H_3_O^+^ at the CL interface is thus not in contradiction with the results of Sparr et al. (26). What has been neglected is the fact that H_3_O^+^ adsorption may cause drastic changes without covalent binding to CL. We are quite aware of the potentially far-reaching biological implications of this finding; however, they are beyond the scope of this study.

**Figure 8.**
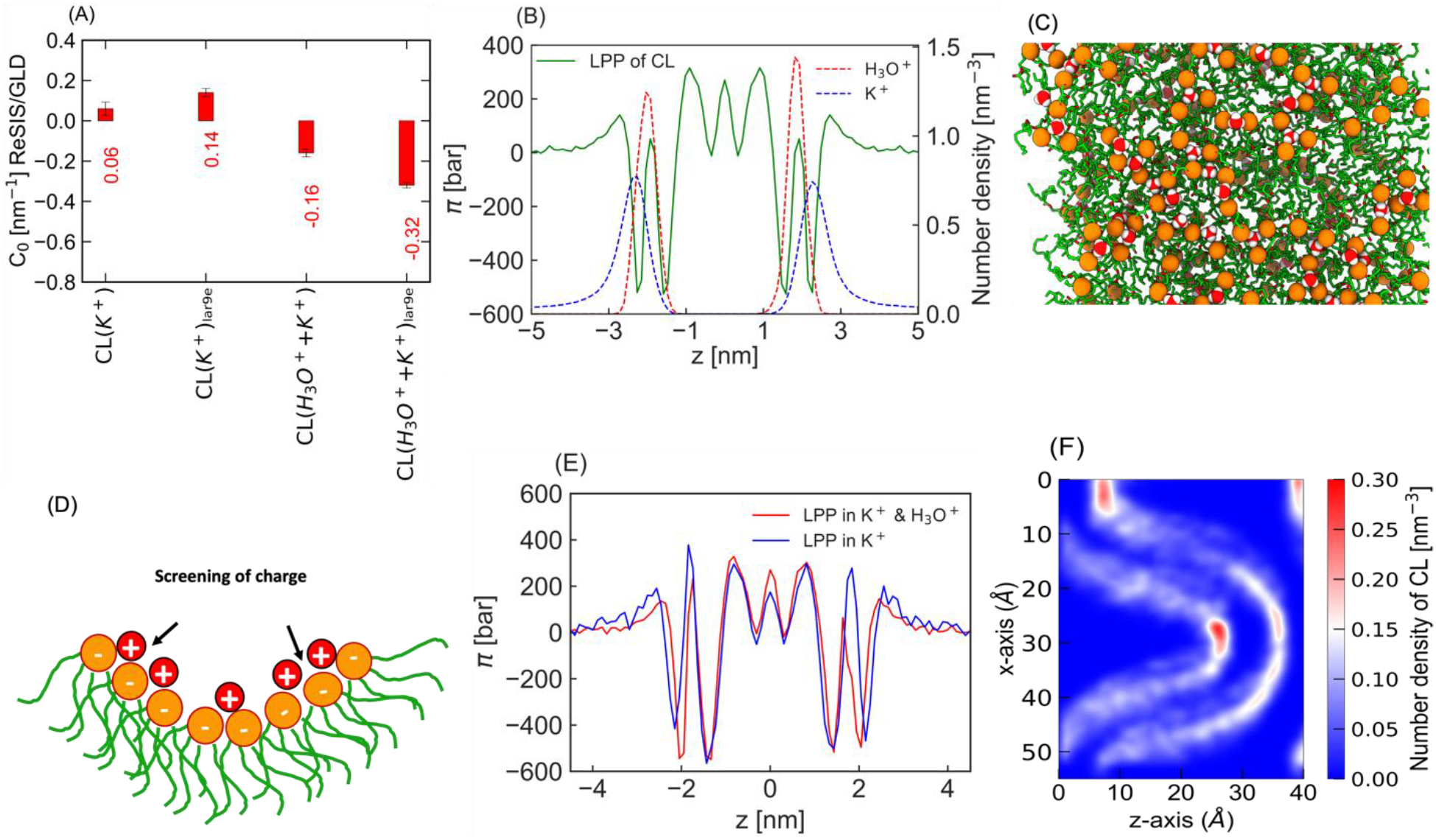
Spontaneous curvature of CL and lateral pressure profile of CL in the presence of K^+^ + H_3_O^+^ ions. (A) Spontaneous curvature (C_0_ value in nm^−1^) of CL (OPLS-AA) in the presence of K^+^ and K^+^ + H_3_O^+^ ions, respectively. (B) Lateral pressure profile of CL with respect to the K^+^ and H_3_O^+^ ion number density. (C) Top view of a frame from a 200 ns simulation trajectory where H_3_O^+^ is adsorbed between the CL head groups, (D) Cartoon representation of CL clustering in the presence of H_3_O^+^

We visualized the trajectory of our simulation, and we found that H_3_O^+^ ions are bridging head groups of CL via hydrogen bonds. Figure 8 D shows the top view of the CL bulk system where the H_3_O^+^ ion is trapped between the head group. Here, the H_3_O^+^ ion is screening the repulsion between two negatively charged head groups of CL. As the repulsion between these two head groups is screened, the CLs are clustered.

The binding of H_3_O^+^ ion to CL should affect the pressure profile of the lipid bilayer thus we have compared the profiles of CL in the presence and the absence of H_3_O^+^ ion. This difference in the lateral pressure profiles of CL is shown in Fig. 8E. The data suggest that H_3_O^+^ ion binding leads to a decline in the first moment of the pressure profile when compared to K^+^ ions. Apart from the significant differences in the headgroup region, the counterion concentration in bulk is also drastically affected, as can be seen in Fig. 8B. Indeed, the spontaneous curvature of CL turns drastically negative upon coordinating H_3_O^+^.

Pressure profiles of bilayers in the lamellar phase only allow the computation of the bending free energy to linear order. The complex interplay of ionic distribution and CL curvature suggests that such an extrapolation might not be permissible. Therefore, we have simulated CL in a strongly curved geometry with K^+^ counterions. The results are shown in Fig. 8F. Results here are interesting, because they show the enrichment of CL at strongly negatively curved parts of the system. But on the other hand, the positively curved parts of the system also display enrichment of CL. We explain this phenomenon again using the counterion density. At the strongly negatively curved interface, the counterion distribution is closer to the membrane due to the lack of space for dissociation as well as due to the reduced permittivity of the adjacent bilayers. Whereas, at the positively curved interfaces, there is a large gap for counterions to distribute. These findings show that CL curvature is not a unique number but is shaped by the counterion distribution, its geometry and specific interactions. It is, therefore, no longer sufficient to speak of a “negative curvature” of CL. Instead, one must consider the CL counterion system and its geometry.

## Conclusion

We have constructed mitochondrial membrane models based on available data for mammals and *Drosophila*. These simple models can be made of commercially available lipids and may guide experimental research. The models are based on simplified membrane compositions and various simplifying estimates, but they reflect the significantly different membrane compositions across species. The difference becomes even more pronounced when considering lipid membrane asymmetry. Simulation of these mitochondrial membrane models with all-atom molecular dynamics allowed us to extract their bulk mechanical properties. We find these properties to be remarkably similar across species, even when including asymmetry. In addition, we find these properties to be in good agreement with common extrapolation schemes. This indicates that the previous approaches to modeling complex membrane compositions agree with detailed simulations.

The main exception in cardiolipin, whose elastic properties, in particular its curvature, are heavily affected by the counterion distribution as exemplified by the adsorption of H_3_O^+^. This finding illustrates that it is not permissible to assign a constant spontaneous curvature to charged lipids without considering the detailed boundary conditions and ion-specific effects. We believe that this insight will also be crucial when explaining curvature sorting observations of cardiolipin and gives a hint towards the role of CL inside proton-gradient generating organelles. Furthermore, we examined the effects of forcefields and CL oxidation and provided elastic parameters for single lipids.

## Supporting Material

Supporting material includes detailed IMM and OMM compositions of different species, an asymmetric lipid composition in the matrix and IMS side of IMM for mammals, and tilt moduli of the bulk lipid bilayer systems and the model mitochondrial membranes. Details about the buckled bilayer composition, asymmetric membrane preparation, area per lipid, total area of each leaflet and bilayer thickness are also included.

## Acknowledgements

This work was funded by the Charles University PRIMUS Grant PRIMUS/20/SCI/015 (C.A., H.A. and S.K.). H. A. was also funded by the Charles University grant UNCE/SCI/023. We thank Prof. Luke H. Chao for bringing to our attention the challenges of mitochondrial membrane morphology and Dr. Piotr Jurkiewicz for discussing CL biophysics with us.

