## Supporting Material for "Mitochondrial Membranes: Model Lipid Compositions, Material Properties and the Changing Curvature of Cardiolipin"

### 1) Lipid headgroup composition of IMM and OMM for different species

Here, we calculated the ratio of the lipids between IMM and OMM. From Table S1, we suggest that the ratio of headgroups in IMM and OMM remains almost the same across different species. The headgroup data for Table S1 is obtained from Horvath et al.(1), Table 4.

**TABLE S1: Lipid headgroup composition of IMM and OMM for different species**

| Species | Lipid | (IMM/OMM) | $N_{IMM(Lipid)}$ | $N_{OMM(Lipid)}$ |
|---|---|---|---|---|
| Mammalian cells (Rat liver) | PC | (40/54)=0.740 | 40 | 54 |
| Plant cells (Cauliflower) | PC | (42/47)= 0.893 | 42 | 47 |
| Yeast (Saccharomyces cerevisiae) | PC | (38/46)= 0.826 | 38 | 46 |
| Mammalian cells (Rat liver) | PE | (34/29)=1.172 | 34 | 29 |
| Plant cells (Cauliflower) | PE | (38/27)=1.407 | 38 | 27 |
| Yeast (Saccharomyces cerevisiae) | PE | (24/33)=0.727 | 24 | 33 |
| Mammalian cells (Rat liver) | PI | (5/13)=0.384 | 5 | 13 |
| Plant cells (Cauliflower) | PI | (5/23)=0.217 | 5 | 23 |
| Yeast (Saccharomyces cerevisiae) | PI | (10/16)=0.625 | 10 | 16 |
| Mammalian cells (Rat liver) | PS | (3/2)=1.5 | 3 | 2 |
| Plant cells (Cauliflower) | PS | - | - | - |
| Yeast (Saccharomyces cerevisiae) | PS | (4/1)=4 | 4 | 1 |
| Mammalian cells (Rat liver) | CL | (18/1)=18 | 18 | 1 |
| Plant cells (Cauliflower) | CL | (15/3)=5 | 15 | 3 |
| Yeast (Saccharomyces cerevisiae) | CL | (16/6)=2.666 | 16 | 6 |

### 2) Lipid distribution in IMM and OMM for mammals

The distribution of phospholipids in mammalian IMM and OMM based on acyl chain composition and head group of mammals is presented in Table S2 and Table S3, respectively. The head group data is based on Table S1 (rat liver data). The acyl chain composition presented in the current article closely resembles with the literature data obtained from Ardail et al. (2), Table II.

We chose to increase the amount of monounsaturated lipids 18:1 with respect to 18:0, because we wanted to examine the physical properties of the single component lipids under conditions consistent with the membrane simulation. Fully unsaturated lipids with long tails are not fluid at room temperature, and hence are incompatible with the standard conditions of our simulations.

**TABLE S2. Mammalian IMM composition including headroup and acyl chain information**

| Lipids | PE | PC | CL | PI | PS | Total acyl variant | % of acyl chain | Acyl chain composition(2) | Simplify % |
|---|---|---|---|---|---|---|---|---|---|
| 16:0/18:1 | 34 | 36 | 0 | 0 | 0 | 70 | 31.777 (16:0) | 29 % (16:0) | 45 |
| 16:1/16:1 | 0 | 4 | 0 | 0 | 0 | 4 | 03.389 (16:1) | 02 % (16:1) | 2 |
| 18:1/18:1 | 0 | 0 | 0 | 0 | 0 | 0 | 29.661 (18:1) | 17% (18:1) | 17 |
| 18:2/18:2 | 0 | 0 | 18 | 0 | 0 | 36 | 33.898 (18:2) | 24% (18:2) | 36 |
| 16:0/18:2 | 0 | 0 | 0 | 5 | 0 | 5 | | | |
| 18:0/18:2 | 0 | 0 | 0 | 0 | 3 | 3 | 01.271 (18:0) | 16% (18:0) | |
| Total | 34 | 40 | 18 | 5 | 3 | 118 | 100 | 12%(20:4) | 100 |

**TABLE S3. Mammalian OMM composition including headroup and acyl chain information**

| Lipids | PE | PC | CL | PI | PS | Total acyl variant | % of acyl chain | Acyl chain composition(2) | Simplify % |
|---|---|---|---|---|---|---|---|---|---|
| 16:0/18:1 | 25 | 50 | 0 | 0 | 0 | 75 | 42.64(16:0) | 29 % (16:0) | 45 |
| 16:1/16:1 | 4 | 4 | 0 | 0 | 0 | 8 | 7.8431(16:1) | 2 % (16:1) | 2 |
| 18:1/18:1 | 0 | 0 | 0 | 0 | 0 | 0 | 36.764(18:1) | 17% (18:1) | 17 |
| 18:2/18:2 | 0 | 0 | 2 | 0 | 0 | 4 | 11.274(18:2) | 24% (18:2) | 36 |
| 16:0/18:2 | 0 | 0 | 0 | 12 | 0 | 12 | | | |
| 18:0/18:2 | 0 | 0 | 0 | 0 | 3 | 3 | 01.470 (18:0) | 16% (18:0) | |
| Total | 29 | 54 | 2 | 12 | 3 | 102 | 100 | 12% (20:4) | 100 |

In Table S4, the lipid composition of IMM and OMM based on the data obtained from Horvath et al.(1), Table 1.

**TABLE S4. Lipid headgroup composition of IMM and OMM for mammals (rat liver)**

| Lipid | % of total phospholipid(1) | (IMM/OMM)(1) | $N_{IMM(Lipid)}$ | $N_{OMM(Lipid)}$ | Avg (Lipid) |
|---|---|---|---|---|---|
| PC | 44 | (40/54) = 0.740 | 40 | 54 | 42.8 |
| PE | 34 | (34/29) = 1.172 | 34 | 29 | 33 |
| PI | 5 | (5/13) = 0.3846 | 5 | 13 | 6.8 |
| CL | 14 | (18/1.0) = 18 | 18 | 1 | 14.8 |
| PS | 3 | (3/3) =1.00 | 3 | 3 | 3 |

#### 3) Lipid composition of IMM and OMM for *Drosophila*

The distribution of lipids in the IMM and OMM of Drosophila is presented in Table S5. The lipid head group data is obtained from Acehan et al.(3), Figure 3. However, the lipid ratio in IMM and OMM is calculated according to Table S4.

**TABLE S5. Estimated Lipid headgroup composition of IMM and OMM in Drosophila**

| Lipid | % of total phospholipid(3) | (IMM/OMM)(1) | $N_{IMM(Lipid)}$ | $N_{OMM(Lipid)}$ | Avg (Lipid) |
|---|---|---|---|---|---|
| PC | 23 | (20/27) = 0.74 | 20 | 27 | 21.4 |
| PE | 48 | (53/45) = 1.17 | 53 | 45 | 51.4 |
| PI | 13 | (10/25) = 0.4 | 10 | 25 | 13 |
| CL | 6 | (8/1.0) = 8 | 8 | 1 | 6.4 |
| PS | 10 | (9/6) = 1.5 | 9 | 6 | 7.8 |

The distribution of phospholipids in IMM and OMM including acyl chain composition along the head group of *Drosophila* is presented in Table S6 and Table S7. The acyl chain composition is in good agreement with the data from Dubessay et al.(4), Table 3.

**TABLE S6. Model Lipid distribution of Drosophila IMM**

| Lipids | PE | PC | CL | PI | PS | Total acyl variant | % of acyl chain | Acyl chain composition(4) | Simplify % |
|---|---|---|---|---|---|---|---|---|---|
| 16:0/18:1 | 12 | 6 | 0 | 0 | 0 | 18 | 12.962(16:0) | 12 % (16:0) | 16 |
| 16:1/16:1 | 24 | 15 | 0 | 0 | 0 | 39 | 36.111(16:1) | 34 % (16:1) | 34 |
| 18:1/18:1 | 12 | 4 | 0 | 0 | 0 | 16 | 23.148(18:1) | 24% (18:1) | 24 |
| 18:2/18:2 | 0 | 0 | 8 | 0 | 0 | 16 | 23.611(18:2) | 26% (18:2) | 26 |
| 16:0/18:2 | 0 | 0 | 0 | 10 | 0 | 10 | | | |
| 18:0/18:2 | 0 | 0 | 0 | 0 | 9 | 9 | 04.166 (18:0) | 01% (18:0) | |
| Total | 48 | 25 | 8 | 10 | 9 | 108 | 100 | 3 % (14:0) | |

**TABLE S7. Model lipid distribution of the OMM of Drosophila**

| Lipids | PE | PC | CL | PI | PS | Total acyl variant | % of acyl chain | Acyl chain composition(4) | Simplify % |
|---|---|---|---|---|---|---|---|---|---|
| 16:0/18:1 | 3 | 7 | 0 | 0 | 0 | 10 | 12.745(16:0) | 12% (16:0) | 16 |
| 16:1/16:1 | 29 | 10 | 0 | 0 | 0 | 39 | 38.235 (16:1) | 34% (16:1) | 34 |
| 18:1/18:1 | 16 | 8 | 0 | 0 | 0 | 24 | 28.431 (18:1) | 24% (18:1) | 24 |
| 18:2/18:2 | 0 | 0 | 2 | 0 | 0 | 4 | 16.176 (18:2) | 26% (18:2) | 26 |
| 16:0/18:2 | 0 | 0 | 0 | 16 | 0 | 16 | | | |
| 18:0/18:2 | 0 | 0 | 0 | 0 | 9 | 9 | 04.411 (18:0) | 01% (18:0) | |
| Total | 48 | 25 | 2 | 16 | 9 | 102 | 100 | 14:0 (3 %) | |

### 4) Asymmetric Lipid composition of the matrix and IMS side of mammal IMM

The asymmetric lipid distribution of the matrix side and IMS of mammal IMM is presented in Table S8. The distribution of phospholipids follows the composition of IMM for mammals, with the aim of an equal area for both leaflets. We incorporated asymmetrical headgroup information from (1).

**Table S8. Asymmetric lipid distribution in the IMS and Matrix side of the mammal IMM**

| IMM | Matrix side | | | | | IMS side | | | | | | |
|---|---|---|---|---|---|---|---|---|---|---|---|---|
| Lipids | PE | PC | CL | PI | PS | PE | PC | CL | PI | PS | Total | Tail % |
| 16:0/18:1 | 34 | 16 | 0 | 0 | 0 | 34 | 56 | 0 | 0 | 0 | 140 | 31.779 (16:0) |
| 16:1/16:1 | 0 | 6 | 0 | 0 | 0 | 0 | 2 | 0 | 0 | 0 | 8 | 03.389 (16:1) |
| 18:1/18:1 | 0 | 0 | 0 | 0 | 0 | 0 | 0 | 0 | 0 | 0 | 0 | 29.661 (18:1) |
| 18:2/18:2 | 0 | 0 | 27 | 0 | 0 | 0 | 0 | 9 | 0 | 0 | 72 | 33.898 (18:2) |
| 16:0/18:2 | 0 | 0 | 0 | 8 | 0 | 0 | 0 | 0 | 2 | 0 | 10 | |
| 18:0/18:2 | 0 | 0 | 0 | 0 | 0 | 0 | 0 | 0 | 0 | 6 | 6 | 01.271 (18:0) |
| Total | 34 | 22 | 27 | 8 | 0 | 34 | 58 | 9 | 2 | 6 | 236 | 100 |

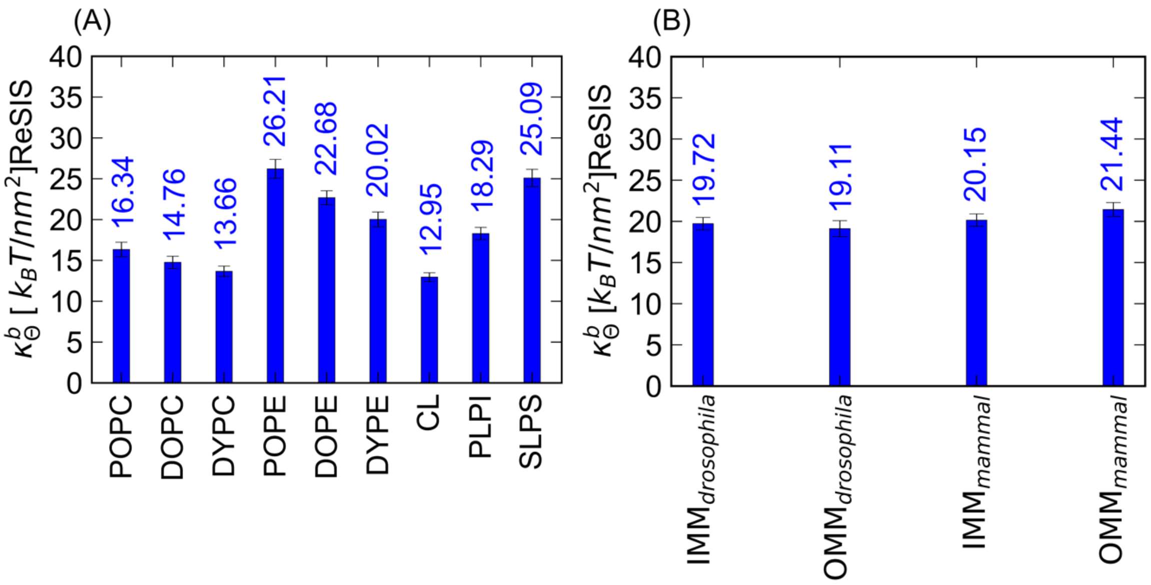

**Figure S1. Tilt moduli of bulk and model IMM and OMM for different species.** Tilt moduli ($\kappa_\theta^b$) (in kT/nm$^2$ unit) of (A) the bulk lipid bilayer system and (B) the model mitochondrial membranes of both species.

**TABLE S9. Simulation System Compositions**

| Name | Lipids | Water | K+ | Cl- | T (K) |
|---|---|---|---|---|---|
| **DOPE** | DOPE 200 | 11529 | | | 303.15 |
| **DYPE** | DYPE 200 | 11231 | | | 303.15 |
| **POPE** | POPE 200 | 10610 | | | 303.15 |
| **DOPC** | DOPC 200 | 12535 | | | 303.15 |
| **DYPC** | DYPC 200 | 12021 | | | 303.15 |
| **POPC** | POPC 128 | 4863 | | | 303.15 |
| **PLPI** | PI 200 | 10821 | 227 | 27 | 303.15 |
| **SLPS** | PS 200 | 10771 | 227 | 27 | 303.15 |
| **CL (CHARMM) (small box)** | CL 200 | 18125 | 448 | 48 | 303.15 |
| **CL (CHARMM) (large box)** | CL 200 | 86330 | 640 | 240 | 303.15 |
| **CL (OPLS-AA)** | CL 200 | 18126 | 448 | 48 | 303.15 |
| **CLox** | CL 200 | 18126 | 448 | 48 | 303.15 |
| **IMM $_{\mathbf{drosophila}}$** | Lipids 200* | 12554 | 102 | 32 | 303.15 |
| **OMM $_{\mathbf{drosophila}}$** | Lipids 200* | 11313 | 85 | 29 | 303.15 |
| **IMM $_{\mathbf{mammal}}$** | Lipids 200* | 13923 | 124 | 36 | 303.15 |
| **OMM $_{\mathbf{mammal}}$** | Lipids 200* | 11750 | 68 | 30 | 303.15 |
| $IMM_{asymm}$ | Lipids 200* | 12000 | 115 | 27 | 303.15 |
| $\boldsymbol{IMS_b}$ | Lipids 218* | 13080 | 82 | 30 | 303.15 |
| $\boldsymbol{Matrix_b}$ | Lipids 182* | 10920 | 148 | 24 | 303.15 |
| **Wave IMM (Mammal)** | Lipids 800* | 67520 | 526 | 174 | 303.15 |

### 1. Additional Simulation Details

#### 1.1. Details about buckled bilayer system preparation

We built the buckled system by assembling four equilibrated IMM patches into a box of 8.65 nm width and 4x 8.65 nm length. We then set the compressibility in width to zero and applied a pressure difference between length and height (direction of the water) until the system shrank to a length of 21.99630 nm. In the next step, we also set compressibility in the length direction to zero and equilibrated the system using semi-isotropic pressure coupling at 1 bar. The system was simulated for a total of 980 ns. We used the same setups in the Computational Details and in the main manuscript, with the exception of the CSVR thermostat (5). The first 200 ns were not used for sampling.

#### 1.2. Details about the Asymmetric membrane preparation

For asymmetric membrane modeling, we prepared two different bilayers that contain symmetrically distributed phospholipids. The two bilayers $Matrix_b$ and the $IMS_b$ are prepared according to the compositions of phospholipids mentioned in Table S8. After that, both of the bilayers were equilibrated in an NPT ensemble. For equilibration, we used the same method as mentioned previously (in the method section). After that, we combined the monolayer compositions of each of the bilayer systems. In this way, we obtained the final asymmetric membrane ($IMM_{asymm}$). Thus, in our model $IMM_{asymm}$both of the layers had different lipid compositions. Furthermore, the $IMM_{asymm}$ was equilibrated for 100 ns and then sampled for 200 ns in an NpT ensemble (Details provided in the method section). The leaflets of the $IMM_{asymm}$are denoted as $Matrix_a$ and $IMS_a$, respectively. More specifically, $Matrix_a$ is the inner leaflet of the IMM, which faces the mitochondrial matrix and $IMS_a$ is outer leaflet represents part of the membrane facing the intramembrane space.

**Table S10. Area per lipid and bilayer thickness of bulk lipid bilayers for Drosophila and Mammals**

| Lipid type | Area Per Lipid [APL] (Å$^2$) | Bilayer thickness (P-P) (Å) |
|---|---|---|
| DOPC | 68.02 ± 0.08 | 38.48 |
| DYPC | 68.20 ± 0.20 | 35.13 |
| POPC | 65.80±1.40 | 39.21 |
| $CL_{CHARMM}$ | 134.29 ± 0.20 | 37.26 |
| $CL_{OPLS-AA}$ | 132.33±0.12 | 38.11 |
| $CL_{ox\ (OPLS-AA)}$ | 140.21±0.67 | 36.12 |
| DOPE | 61.64 ± 0.16 | 40.98 |
| DYPE | 61.85 ± 0.11 | 37.18 |
| POPE | 58.97± 0.06 | 41.34 |
| PLPI | 63.90 ± 0.17 | 38.63 |
| SLPS | 59.93 ± 0.12 | 42.17 |
| $IMM_{drosophila}$ | 67.26 ± 0.99 | 38.95 |
| $OMM_{drosophila}$ | 63.14 ± 0.98 | 38.59 |
| $IMM_{mammal}$ | 73.13 ± 1.06 | 39.75 |
| $OMM_{mammal}$ | 62.80 ± 0.14 | 39.80 |
| $Matrix_b$ | 74.71 ± 1.08 | 25.21 |
| $IMS_b$ | 72.10 ± 1.02 | 22.06 |
| $IMM_{asymm}$ | 73.35 ± 0.97 | 48.73 |

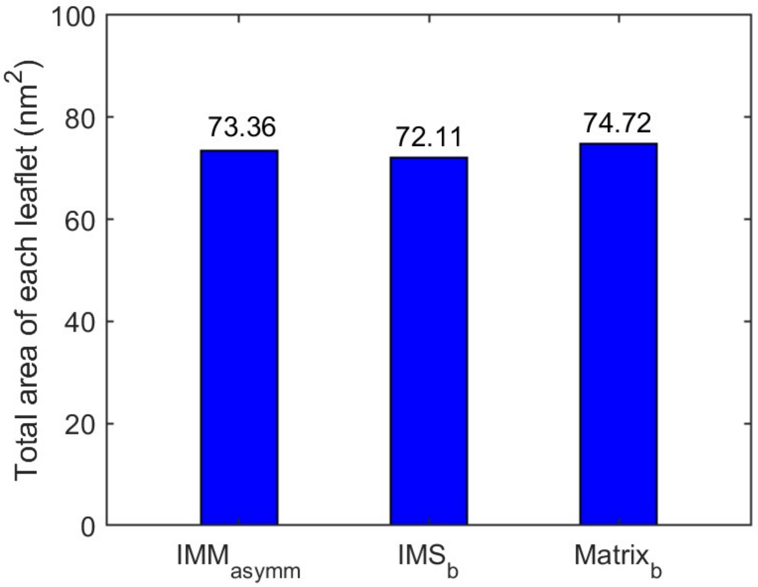

**Figure S2. Total membrane leaflet area of the asymmetrical mammalian IMM simulation in comparision with symmetrical simulations of its IMS and Matrix sides.**
